# Explainability methods from machine learning detect important drugs’ atoms in drug-target interactions

**DOI:** 10.64898/2026.01.02.697342

**Authors:** Mrinal Mahindran, Qingyuan (Chingyuen) Liu, Vishak Madhwaraj Kadambalithaya, Olga V Kalinina

## Abstract

Predicting drug-target interactions (DTI) with graph neural networks (GNNs) is hindered by their lack of interpretability. To address this, we benchmark four explainable artificial intelligence (XAI) attribution methods on GNN models trained for kinase and GPCR targets. We assess the methods’ consistency through atom-level intersection-over-union and validate their biological relevance by mapping attributed atoms to 3D protein-ligand structures. While consistency across methods was modest, consensus attributions were highly enriched for atoms directly contacting the binding pocket—up to 76% within 2 Å in the kinase-inhibitor complexes. Notably, these attributed atoms were frequently found contacting experimentally important regulatory residues, such as those in the DFG motif. This indicates that XAI methods, despite their disagreements, can identify chemically meaningful ligand features, providing a foundation for developing more interpretable GNNs in drug discovery.

## Introduction

In drug discovery, identifying viable drug candidates from vast library of molecules presents a profoundly complex challenge. ^1,2^ Computational drug-target interaction (DTI) prediction methods have become essential tools to bridge the gap between theoretical exploration of the chemical space and resource-intensive experimental validation (e.g., high-throughput screening).^3,4^ Currently, deep learning approaches represent the most widely adopted methods among computational DTI prediction techniques, demonstrating significant utility for enhanced binding affinity prediction, off-target identification, and drug repurposing, among other critical applications in modern-day drug discovery, especially the early stages of drug discovery.^5–7^

Recent advances in deep learning, especially graph neural networks (GNNs), have developed DTI predictions by representing proteins and ligands as graphs, capturing complex chemical and topological dependencies through message-passing frameworks.^8–14^ Among the corresponding architectures, GNNs have demonstrated superior ability in modeling molecular interactions despite higher computational demands compared to classical machine learning or deep learning models. ^15,16^ Their strength lies in approximating highly non-linear structure-activity relationships directly from raw graph representations of chemical structures, ^17,18^ potentially superseding classic hand-crafted molecular fingerprints.^17^ Nevertheless, adoption of GNNs in drug discovery remains limited due to their “black-box” nature and the inability to provide chemically intuitive explanations for predictions. ^2,19^ This interpretability gap is further compounded by risks such as the Clever Hans effect: producing correct answers for the wrong reasons, ^20^ and overconfident erroneous outputs.^21^ Explainable artificial intelligence (XAI) methods aim to address these issues by making GNN decision processes transparent^22–26^ – revealing which sub-structural motifs in the ligand and which residue interactions in the protein drive the predicted binding – thus bridging machine reasoning with chemical intuition. ^2,27–30^

Multiple existing studies have addressed these issues, primarily through the visualization of attention weights.^31,32^ While the application of attention mechanisms is straight-forward and effective, it remains limited to attention- or transformer-based GNN architectures. Furthermore, such mechanisms typically capture only local vertex neighborhoods (often termed masked attention),^33,34^ failing to elucidate global relationships between atoms.^35^ In contrast, gradient-based XAI methods offer greater universality across diverse GNN architectures and have demonstrated considerable promise.^35^ To our knowledge, however, no comprehensive benchmark exists for evaluating gradient-based attribution methods specifically on DTI GNN models. To address this gap, our study implements and assesses four attribution-based XAI techniques: input × gradient,^36^ guided backpropagation,^37^ integrated gradients,^38^ and gradients SHAP.^39^ We apply these to GNN-based DTI models trained on two curated datasets from the ligands of two distinct families of proteins: the G protein-coupled receptor (GPCR) ligands^40^ and kinase inhibitors.^4,41^ We trained the models in four different data splitting regimes: random, cold-drug, cold-target, scaffold-along with an additional biological-relevance split, and then applied the attribution methods to the best-performing model for analysis. We introduce complementary assessments: first, quantifying the explanation agreement through intersection-over-union (IoU) of attributed atoms; and second, validating biological relevance by mapping consensus-attributed atoms against known binding site residues and pharmacophoric features. This work presents a systematic framework for benchmarking the consistency of attribution and biological plausibility in GNN explanations for DTI. We provide our evaluation code and consensus analysis pipelines publicly to enable reproducibility and accelerate the development of trustworthy models at https://github.com/kalininalab/LigandXai.

## Methods

### Datasets and Preprocessing

We evaluated our models on two widely used bioactivity datasets: the KIBA dataset of kinase inhibitors^4^ and the GLASS dataset of the association of GPCR-ligands.^40^ The KIBA and GLASS datasets represent two critical resources in drug discovery, each focusing on a therapeutically pivotal protein family.

The KIBA dataset focuses on kinases, representing a critical protein family that, among other functions, regulates signaling pathways in eukaryotic cells,^42^ making them prime drug targets in oncology and immunology.^42^ To describe interactions between kinases and the corresponding inhibitors, the KIBA score integrates three binding metrics (K_*i*_, K_*d*_, IC_50_) into a unified bioactivity value, enabling robust prediction of compound efficacy against kinase subfamilies (e.g., tyrosine kinases). ^4^ The dataset contains 246,088 KIBA scores across 52,498 chemical compounds and 467 kinase targets. The lower KIBA score indicates a higher binding affinity between the kinases and the drugs. For training our models, we removed any drug or target with fewer than 10 total recorded values in the dataset. KIBA scores were then negated and shifted to set the minimum value to zero, preserving bioactivity rankings while ensuring a non-negative regression target.

The GLASS dataset focuses on G protein-coupled receptors (GPCRs), a protein family crucial in drug discovery due to their role in mediating diverse physiological processes (e.g., neurotransmission, immune responses) ^43^ and their status as targets for ~35% of FDA-approved drugs.^43^ The GLASS dataset was first cleaned by removing any interactions that did not have a reference to any database, publication, or if their references were broken links, as their validity could not be confirmed. Any interactions with invalid energy scores (recorded values are not a number) were also discarded. Furthermore, because of the compilation of interactions from various datasets, even after removing entries with the same value and references, some duplicate interactions bypassed filters. The difference between such duplicates was the difference in rounding. Such entries were manually checked and removed. The newly cleaned dataset contains 513,246 unique compound–GPCR pairs (681 GPCRs and 277,651 ligands), down from the original 562,871 pairs. We retained only those entries with explicitly reported K_*i*_ values, which were all transformed to their corresponding pK_*i*_ values and scaled by a factor of 10^−9^ for stabilizing model training^5^ using *pK*_*i*_ = − log_10_(*K*_*i*_ × 10^−9^).

For both datasets, SMILES strings were processed using RDKit^44^ to generate molecular structures, which were further canonicalized and sanitized to ensure consistent atom ordering. Molecular graphs were produced with PyTorch Geometric.^45^ Compounds with more than 150 heavy atoms were excluded to reduce computational overhead, and data points with missing SMILES or affinity values were discarded. The remaining number of data points is summarized in Table 1

**Table 1:** Summary of datasets and binding affinity scores in the clean datasets after filtering.

| Dataset | Proteins | Ligands | Datapoints | Binding Score |
| --- | --- | --- | --- | --- |
| GLASS | 490 | 70,517 | 136,085 | $\text{pK}_i$ ($\log K_i$) |
| KIBA | 421 | 2,353 | 148,327 | KIBA score (normalized) |

### Graph Neural Networks

Message-passing neural networks (MPNNs), a class of graph neural networks (GNNs), form the foundation of our architecture. In this framework, both drug molecules and proteins are represented as graphs *G* = (*V, E*). The molecular graph representation for each drug is constructed from its SMILES string mapped into an undirected graph *G*_*D*_ = (*V*_*D*_, *E*_*D*_). The nodes are characterized by feature vectors 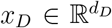 representing one-hot encoded atomic types, while the edges are determined by the molecular bonding topology. Similarly, each protein is represented as a residue-level graph *G*_*P*_ = (*V*_*P*_, *E*_*P*_) derived from its PDB structure. Here, node features 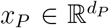 are one-hot encodings of the 20 standard amino acid types, and an edge is created between two residues if the distance between their *α*-carbons (*Cα*) is less than or equal to 8 Å.

The MPNN operates through a series of iterative message-passing layers that update node representations by aggregating information from their local neighborhoods. At each layer *k*, the process involves two main steps. First, in the message aggregation step, each node *v* collects messages from its neighbors 𝒩 (*v*) according to:

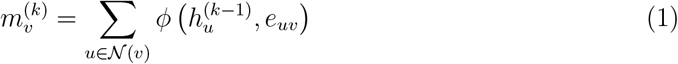

where 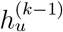 is the feature vector of a neighboring node *u* from the previous layer, *e*_*uv*_ is the feature vector of the edge between them, and *ϕ* is a learnable message function. Second, in the feature update step, the node’s representation is updated by combining its previous state with the aggregated message:

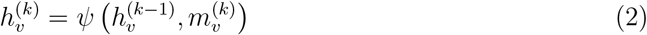

where *ψ* is an update function, typically a neural network with a non-linear activation function such as ReLU. This message-passing scheme is repeated for *K* layers, allowing the model to progressively capture information from larger structural contexts. The final graph-level representations are then used to predict drug-target interactions via downstream neural network architectures. Both the message function *ϕ* and the update function *ψ* contain learnable parameters that are optimized end-to-end during training using backpropagation.

### Model Architecture and Training

The model uses a dual encoder framework, where separate graph neural networks (GNN) encode the ligand and protein graphs. Each encoder processes its respective graph to produce a fixed 256-dimensional vector representation. We experimented with several popular GNN variants including graph isomorphism network (GIN),^46^ GraphSAGE (graph sample and aggregate),^47^ graph attention network (GAT),^33^ transformer-based graph network,^48^ feature-wise linear modulation convolution (FiLMConv),^49^ Chebyshev convolution (ChebConv)^50^ as implemented in the package RINDTI. Each encoder applied multiple message-passing layers followed by mean pooling to obtain a graph-level embedding. The resulting ligand and protein embeddings were then concatenated and passed to a multi-layer perceptron (MLP) for affinity prediction. Models were trained using mean squared error (MSE) loss and optimized with Adam^51^ (learning rate equal to 0.0001) with a batch size of 256, except for attention-based architectures, which were trained with a batch size of 128. Training was conducted for up to 500 epochs with early stopping triggered after 50 epochs without validation loss improvement. All experiments were implemented using PyTorch Geometric, and random seeds were fixed for reproducibility.

### Data Splitting Strategies

To comprehensively evaluate model generalization, we employed four distinct data splitting strategies. The baseline approach was a *random split*, in which the dataset was randomly divided into 80% training, 10% validation, and 10% test sets. This splitting scenario implies that the validation and test sets contain the same drugs and proteins seen during training and thus represents the least challenging scenario. For more rigorous testing, we utilized two *cold* splits. The *cold drug split* ensured that all drug molecules in the validation and test sets were completely unseen during training, a method designed to evaluate how well the model generalizes to novel compounds. The most challenging scenario was the *cold target split*, where proteins in the test set were absent from the training set. This approach assesses the model’s ability to infer binding interactions for unseen biological targets, a crucial capability for applications such as drug repurposing. Additionally, a *scaffold split* was performed, partitioning compounds by their Bemis–Murcko scaffolds to ensure structural dissimilarity between the training, validation, and test sets. Furthermore, to assess the biological plausibility of model attributions, we designed a separate *biological relevance split*, which consisted of moving all drug–target pairs with available protein-ligand complex structures into the test set to enable direct structure-based evaluation. This split was used solely for biological analysis and did not influence training (i.e. hyperparameter tuning etc.) and selection of the GNN models.

### Cross-Dataset Finetuning

To assess the transferability of learned representations, we performed a targeted cross-dataset fine-tuning procedure. For each transfer direction (KIBA → GLASS and GLASS → KIBA), we initialised from the best dual-encoder checkpoint on the source dataset (selected by the lowest test MSE). The encoder parameters, including all message-passing layers of both the drug and protein branches, were frozen. At the same time, the downstream multi-layer perceptron (MLP) prediction head was re-initialised and trained on the training partition of the target dataset. Fine-tuning employed the same optimisation and regularisation settings as the main experiments to ensure comparability.

### Attribution Methods

This section details the feature attribution techniques implemented to interpret the predictions of our graph neural network model for drug-target binding affinity. We evaluate four gradient-based methods: input × gradient,^36^ guided backpropagation,^37^ integrated gradients,^38^ and gradient SHAP.^39^ Each method assigns importance scores, *S*_*v*_ ∈ ℝ to atoms *v* ∈ *V* in the molecular graph, quantifying their contribution to the predicted binding affinity *F* (*G*_*D*_, *G*_*P*_).

#### Attribution Framework

Formally, our trained graph neural network model is represented as a function *F*(*x*) that outputs drug-target binding affinity predictions, where *x* denotes the input feature matrix encoding both the drug molecular graph and protein structure. A feature attribution method computes importance scores for individual input features, specifically the atoms in the drug molecular graph. The attribution function *A* assigns a score *S*_*i*_ to each atom feature *x*(*i*) according to the following relationship:

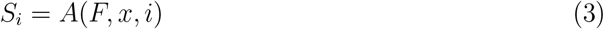

Here, *S*_*i*_ represents the attribution score for the *i*-th atom, quantifying its contribution to the final binding affinity prediction *F*(*x*). The function *A* denotes the specific attribution method used, such as Input × Gradient or Integrated Gradients.

#### Input × Gradient

The input × gradient (I×G) method computes importance scores by calculating the element-wise product of the input feature values and the gradient of the model’s output with respect to those features. The attribution score is formally defined as:

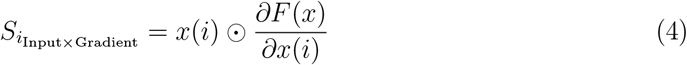

where *x*(*i*) is the input feature of *i*-th atom and 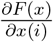 is the gradient of the model output *F*(*x*) with respect to that feature.

#### Guided Backpropagation

The guided backpropagation (GB) method further refines on input × gradient method by only propagating positive gradients through ReLU activation. This filtering mechanism is applied to 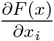, effectively setting gradients to zero if the gradient is negative. This process is intended to reduce noise in the resulting attributions. The final score is then calculated using the filtered gradient:

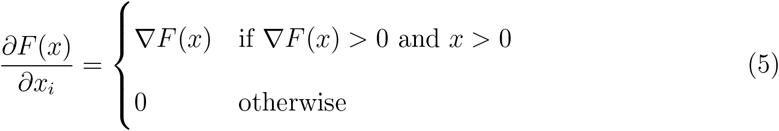

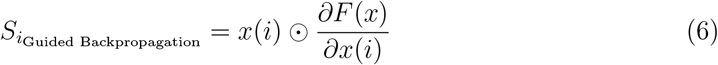

#### Integrated Gradients

The integrated gradients (IG) method computes feature attributions by accumulating gradients along a path from a baseline input *x*′ (typically a zero vector) to the actual input *x*. The method defines a straight-line path *x*(*t*) = *x*′ + *t* · (*x*−*x*′) for *t* ∈ [0, 1]. The attribution is the path integral of the gradients with respect to the input features, scaled by the difference between the input and the baseline:

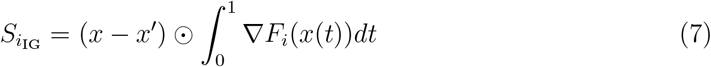

In practice, this integral is approximated numerically by summing the gradients over small intervals along the path.

#### Gradient SHAP

Gradient SHAP extends the integrated gradients framework by improving the choice of baseline. Instead of using a single, fixed baseline, it computes the expectation of the IG attributions over a distribution of baselines, *P* (*x*′). This is typically implemented by sampling multiple baselines {*b*_1_, *b*_2_, …, *b*_*S*_}, often by adding small Gaussian noise to a zero vector, computing the IG attribution for each, and averaging the results:

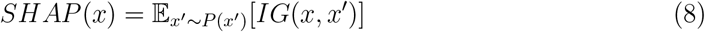

These attribution methods were applied exclusively to the ligand graph, yielding a single attribution score for each atom after processing.

#### Attribution Processing

For atomic-level interpretation, the raw feature-level attribution scores were post-processed. First, for each atom *v*, the scores across all its feature dimensions were aggregated into a single score, *S*_*v*_, by taking their sum. Second, to reduce noise and enhance visual clarity, we effectively discarded any atomic scores where the absolute value |*S*_*v*_| was less than 10^−4^.

### Validation of Biological Relevance

To assess whether the most highly attributed atoms correspond to biologically meaningful regions, we analyzed their spatial proximity to known protein binding pockets using experimentally resolved ligand–protein structures. For the KIBA dataset, we used the KLIFS^52^ database, which provides curated kinase–inhibitor complexes with annotated pocket residues. For GLASS, we retrieved experimentally determined ligand–protein complexes from the RCSB Protein Data Bank (PDB).

We first computed the intersection-over-union (IoU) of the top-k% most attributed atoms for all four explanation methods for each data point (protein–drug interaction), varying *k* across multiple (Table 5). IoU scores were averaged across all data points to obtain a dataset-level measure of agreement. For downstream structural analysis, we focused on the 50% threshold and retained only those data points where (a) the attributed atoms overlapped and (b) the three-dimensional (3D) structural data were available. For each such example, we extracted the consensus-attributed atoms and copied their 3D coordinates from the MOL2 or PDB structures. We subsequently calculated the shortest Euclidean distance from every attributed atom to all atoms in the respective protein binding pocket. Moreover, we measured biological alignment by reporting the proportion of consensus atoms that were within distance cut-offs between 2 Å and 4 Å from the binding pocket residues. This provided a structure-aware and stringent assessment of whether the model’s most prominent predictions overlapped with known interaction sites.

## Results and Discussion

### Model Performance

The model performance of the various GNN-based models was evaluated on both KIBA and GLASS datasets across multiple data splits, including random, cold drug, cold target, and scaffold. In addition, the best-performing model was further evaluated under the biological relevance split, which was reserved solely for downstream structural analysis. Performance was assessed using the MSE between predicted and actual binding affinities. For both datasets, models using GIN for both the protein and drug encoders consistently out-performed other architectures (Table 2). For the KIBA dataset, the random split yielded the best performance, with the GIN_protein_–GIN_drug_ model achieving an MSE of 0.1741. Performance dropped for cold drug and cold protein splits, mirroring the greater challenge of generalizing to new compounds or targets. The scaffold split, which groups molecules based on shared core structures to better test scaffold-level generalization, resulted in intermediate performance (MSE = 0.5859 for KIBA), indicating that while the model retained some generalization ability, it was still challenged by novel molecular scaffolds. For the GLASS dataset, models showed similar trends but with slightly reduced performance overall. The GIN-based model with random splitting achieved an MSE of 0.4766. Notably, while performance again declined for the cold-drug and cold-target splits, the scaffold split exhibited lower MSE than the cold-target split across several architectures, suggesting that generalizing to entirely new targets was more difficult than to unseen molecular scaffolds in this dataset.

**Table 2:** Test MSE⇓ loss comparison across different splits.

| Model | KIBA |  |  |  | GLASS |  |  |  |
| --- | --- | --- | --- | --- | --- | --- | --- | --- |
|  | Random | Cold-drug | Cold-target | Scaffold | Random | Cold-drug | Cold-target | Scaffold |
| GIN | **0.1741** | 0.4607 | 0.4725 | 0.5859 | **0.4766** | 0.5583 | 1.418 | 0.7806 |
| GAT | 0.3228 | 0.5608 | 0.5774 | 0.7577 | 0.7942 | 0.7731 | 1.4573 | 1.036 |
| GraphSAGE | 0.3307 | 0.5936 | 0.7717 | 0.7917 | 0.8559 | 0.8665 | 1.4935 | 1.0273 |
| Transformer-GNN | 0.2864 | 0.5627 | 0.5544 | 0.6236 | 0.699 | 0.7695 | 1.7665 | 1.0678 |
| FiLMConv | 0.1877 | 0.4729 | 0.4866 | 0.672 | 0.5993 | 0.6523 | 1.4901 | 0.8536 |
| ChebConv | 0.4961 | 0.6623 | 0.7891 | 0.6194 | 1.0753 | 1.0374 | 1.5812 | 1.1428 |

We also explored hybrid models where different encoder architectures were used for the protein and drug graphs (Table 3). These models were trained using the random split, since this split consistently yielded the lowest MSE values across both datasets in previous experiments. Even in this comparison, the GIN_protein_–GIN_drug_ model consistently outperformed all the other configurations. To assess robustness, the GIN_protein_–GIN_drug_ model was retrained on the random split across five different seeds. Finally, the selected GIN_protein_–GIN_drug_ model was applied to the biological relevance split to support downstream structural validation, where it achieved an MSE of 0.26 for KIBA and 0.50 for GLASS. Additionally, we explored cross-dataset fine-tuning, where the best model was trained on one dataset and fine-tuned on the other under the random split configuration. The best model (GIN_protein_–GIN_drug_ model) trained on KIBA and fine-tuned on GLASS achieved a test MSE of 0.57, while the same model trained on GLASS and fine-tuned on KIBA achieved a test MSE of 0.24. In both cases, fine-tuned models underperformed the original task-specific models.

**Table 3:** Test Loss (MSE⇓) comparison of different drug and protein encoder combinations on the random split. Values are reported as KIBA / GLASS.

| Drug \ Protein | ChebConv | FiLMConv | GATConv | GINConv | Transformer | GraphSAGE |
| --- | --- | --- | --- | --- | --- | --- |
| ChebConv | 0.496 / 1.075 | 0.503 / 1.200 | 0.462 / 1.049 | 0.493 / 1.021 | 0.471 / 1.041 | 0.487 / 1.053 |
| FiLMConv | 0.228 / 0.608 | 0.187 / 0.599 | 0.223 / 0.605 | 0.182 / 0.571 | 0.194 / 0.591 | 0.225 / 0.571 |
| GATConv | 0.300 / 0.759 | 0.295 / 0.713 | 0.322 / 0.794 | 0.260 / 0.707 | 0.263 / 0.714 | 0.291 / 0.731 |
| GINConv | 0.217 / 0.502 | 0.186 / 0.495 | 0.208 / 0.542 | **0.174 / 0.476** | 0.187 / 0.506 | 0.214 / 0.532 |
| Transformer | 0.301 / 0.765 | 0.281 / 0.764 | 0.283 / 0.820 | 0.269 / 0.703 | 0.286 / 0.699 | 0.283 / 0.867 |
| GraphSAGE | 0.341 / 0.849 | 0.316 / 0.823 | 0.301 / 0.813 | 0.288 / 0.790 | 0.299 / 0.787 | 0.330 / 0.855 |

**Table 4:** Test MSE for the GIN–GIN configuration. Values are mean ± standard error over five seeds.

| Dataset | Drug encoder | Target encoder | Test MSE (mean ± SEM) |
| --- | --- | --- | --- |
| GLASS | GIN | GIN | 0.4893 ± 0.0045 |
| KIBA | GIN | GIN | 0.174 ± 0.0018 |

**Table 5:** Intersection-over-Union (IoU) of important-atom sets at different top-*k* attribution thresholds for KIBA and GLASS.

| Top-$k$ | KIBA | GLASS |
| --- | --- | --- |
| 10% | 0.0140 | 0.0141 |
| 20% | 0.0304 | 0.0333 |
| 30% | 0.0570 | 0.0647 |
| 40% | 0.0972 | 0.1066 |
| 50% | 0.1526 | 0.1636 |

These results indicate that while GNNs are capable of learning expressive representations for drug–target pairs, their ability to generalize is strongly influenced by the target diversity in the dataset and the type of data split used. In particular, it is very difficult to train a well-performing model that would not be protein family-specific. Based on its consistently superior performance across both datasets, the GIN_protein_–GIN_drug_ model was selected for all downstream analyses of explainability and biological relevance.

### Attribution Consistency Across Methods

While basic gradient-based explainability methods such as input × gradient and guided backpropagation compute 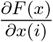, these suffer from gradient saturation^53^ and violate axiomatic properties such as *sensitivity* (nonzero attribution for feature-impacting predictions) and *implementation invariance* (consistent attributions for functionally equivalent models).^38^ We therefore also implemented advanced techniques such as integrated gradients and gradient SHAP. To assess attribution consistency, we computed the intersection-over-union (IoU) for the top 50% most attributed atoms across all four explanation methods in the biological relevance test split, using the best-performing GIN model.

For the GLASS data set, the average IoU score was 0.1636 ± 0.0904 for 30,922 drug-target pairs, while for the KIBA data set, the average IoU score was 0.15 ± 0.0930 for 33,236 drug-target pairs. This reflects a modest level of agreement among the four different attribution methods for both data sets (Figure 1). Only data points with non-zero overlap and available structural data in the test dataset were selected for downstream analysis.

**Figure 1.**
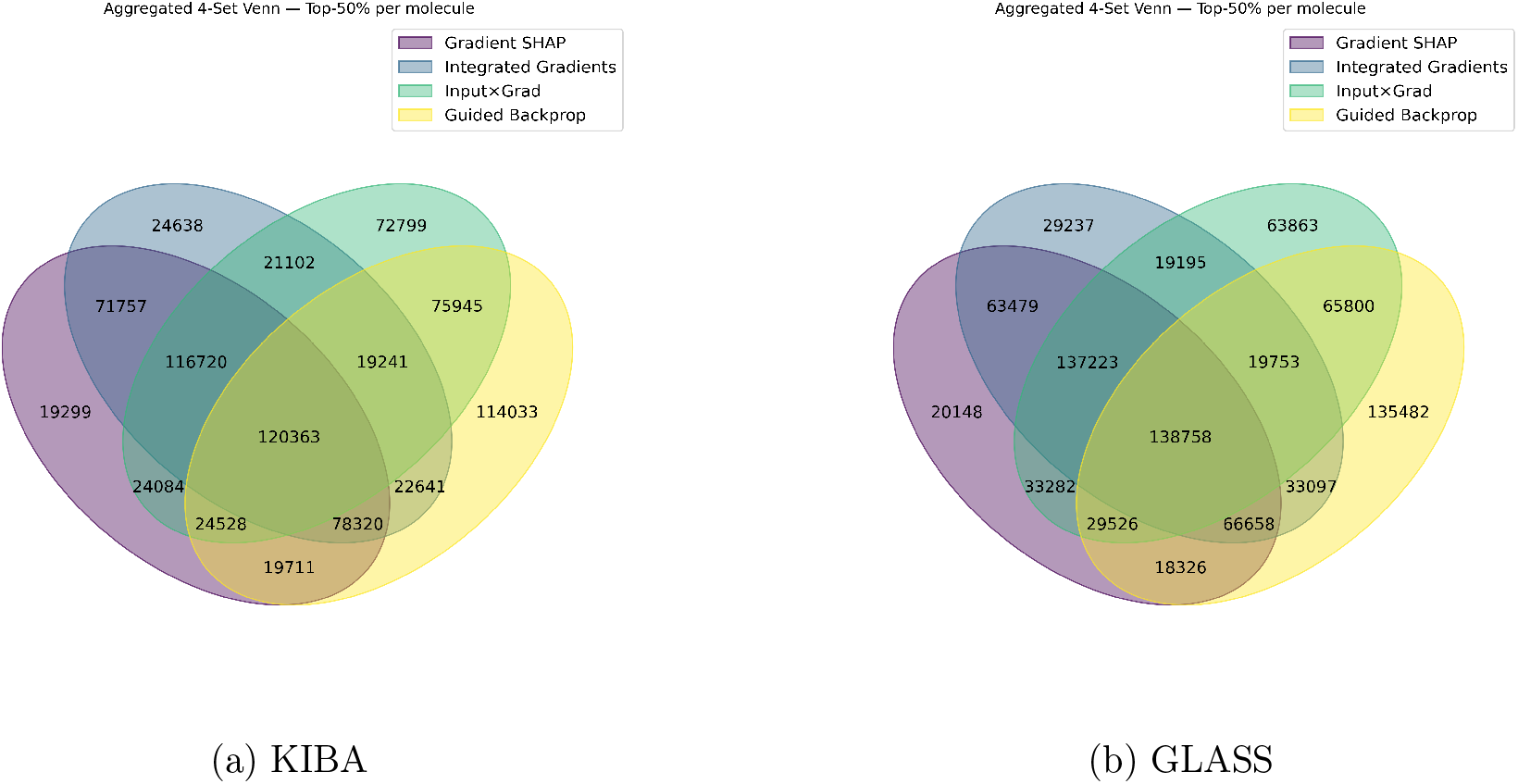
Four-set Venn diagrams for the KIBA (left) and GLASS (right) datasets. The diagrams display the exclusive counts of atoms identified by different subsets of the four explanation methods.

### Biological Relevance of Attributions

To determine whether the attributed atoms correspond to functionally important parts of the small molecules, we quantified their proximity to binding pocket residues in the three-dimensional (3D) structures of experimentally solved complexes of proteins with small molecules (see Supplementary Information). In the KIBA dataset (759 experimentally solved complexes), attributed atoms are highly enriched in a very close proximity to the determined binding pocket annotated from KLIFs: at the 2 Å cutoff, nearly 76% of ligand atoms were marked as important by all four attribution methods. This enrichment declines as the distance grows, reaching 15% at 3 Å and 13% at 4 Å, as the number of ligand atoms within the cutoff increased faster than the number of attributed atoms (Figure 2a,c). In the GLASS dataset (161 experimentally solved complexes), a similar trend was observed: at 2 Å, 33% of ligand atoms were attributed, i.e. marked as important, with enrichment decreasing to 15% at 3 Å and 11% at 4 Å (Figure 2b,d).

**Figure 2.**
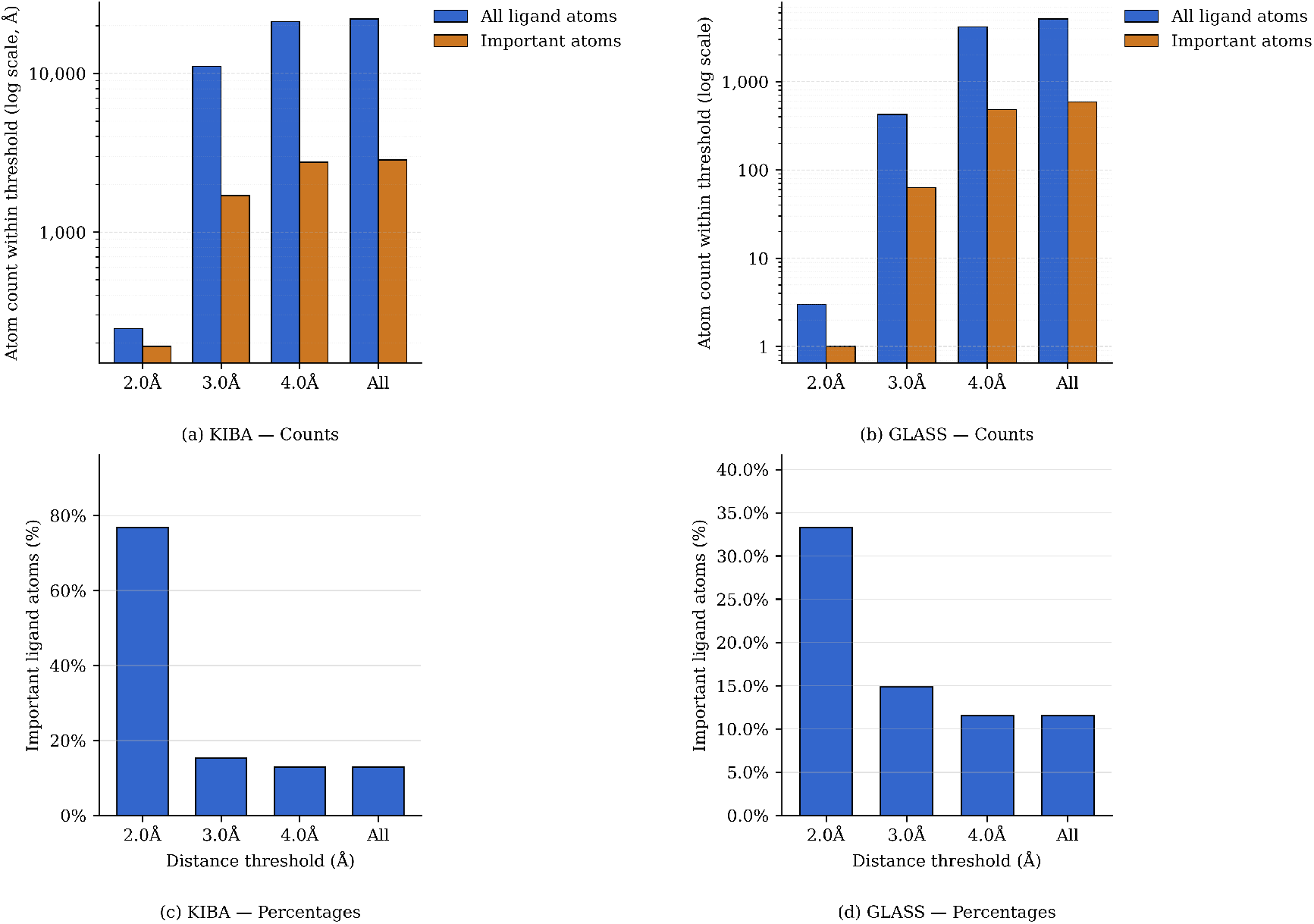
Comparison of atom counts and percentages across distance thresholds for the KIBA and GLASS datasets.

In summary, the consensus attributions, despite modest agreement between the methods, preferentially highlight atoms directly interacting with the protein pocket. This effect is more pronounced in the KIBA dataset, where a larger number of experimentally solved complexes supports strong and reproducible enrichment at direct contacts. In GLASS, the enrichment is weaker but still evident, indicating that consensus attributions capture biologically meaningful ligand substructures even in a smaller and more heterogeneous set of complexes.

For example, in the complex of mitogen-activated protein kinase 14 (MAPK14; UniProt assession Q16539) and the compound with the ChEMBL identifier CHEMBL328242, four of the eight attributed atoms are in direct contact with the protein (Figure 3a), and in the complex of endothelin receptor type B (EDNRB; UniProt assession P24530) and bosentan (ChEMBL identifier CHEMBL957) seven of the seven interact with the protein directly (Figure 3b).

**Figure 3.**
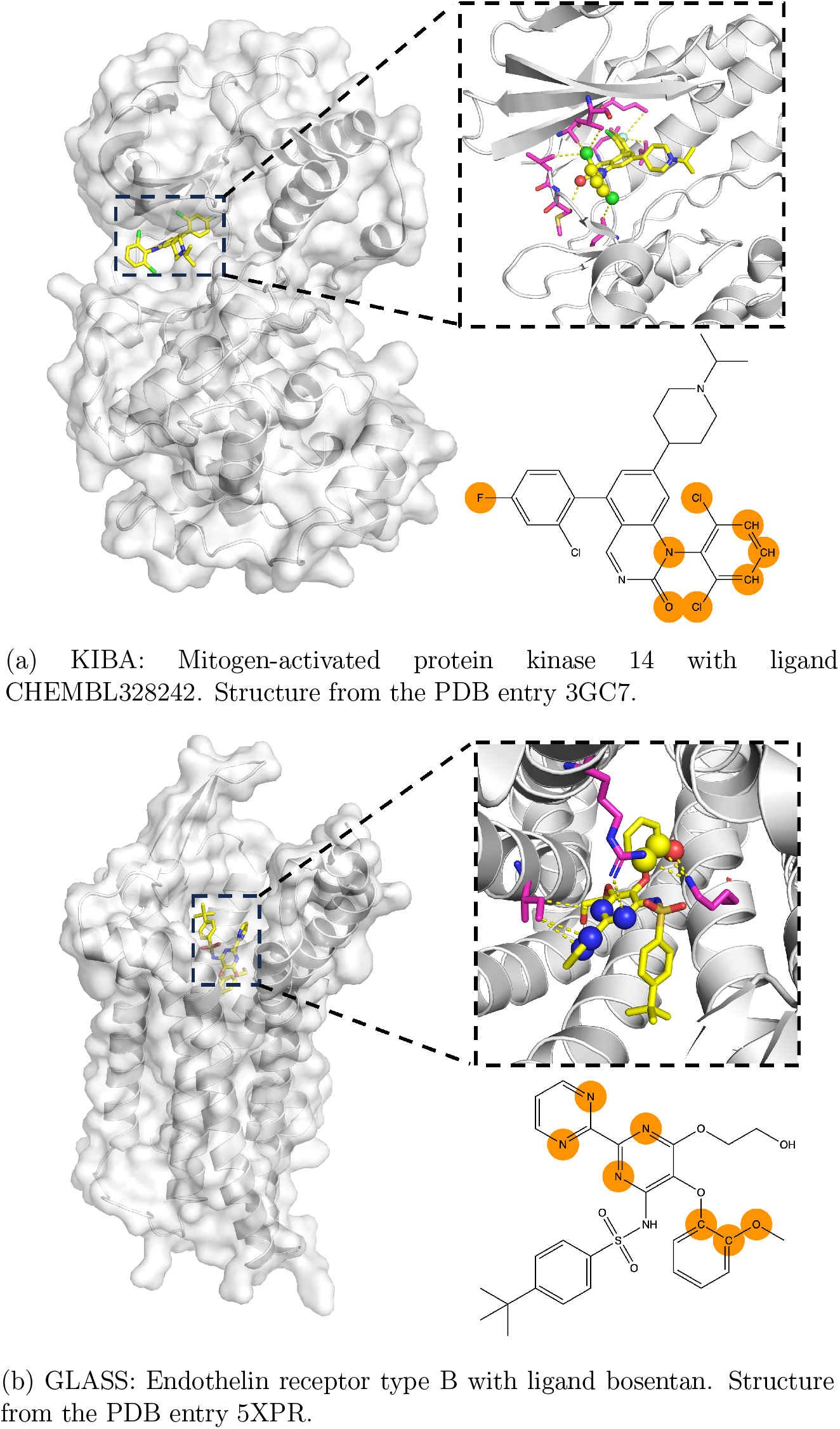
Example complexes from the GLASS and KIBA datasets with consensus-attributed atoms. Attributed atoms are shown as balls in the stick representation of the ligands. Contacting amino acid residues of the protein are shown in sticks, as well, and colored magenta.

For 69 proteins from the KIBA dataset and 43 proteins from the GLASS dataset, several small molecules were observed to interact with the same protein. In these proteins, the attributed atoms of the different interactors were compared to assess consistency across ligands binding to the same protein. Three of the available ligands were chosen for the KIBA dataset and GLASS dataset, respectively. The corresponding protein–ligand complexes were overlaid, and the consensus-attributed atoms were examined to evaluate how well they mapped in three-dimensional space. For the KIBA dataset, the overlay of three ligands with the ChEMBL identifiers CHEMBL2029678, CHEMBL2029688, and CHEMBL2031893 of the hepatocyte growth factor (HGF) receptor (UniProt assession P08581) revealed that the consensus-attributed atoms were consistently located near the DFG motif, a well-established regulatory element of the kinase active site (Figure 4a). Despite differences in their chemical scaffolds, the highlighted atoms converged within this conserved pocket region, suggesting that the explanation methods capture structural determinants that are central to kinase inhibition. The spatial alignment of attributed atoms across different ligands reinforces the view that the model recognizes ligand atoms in the proximity of activity-relevant pocket residues, implying a recognition of biologically meaningful motifs within the kinase binding site. As an example from the GLASS dataset, we analyzed three ligands of the melatonin receptor type 1A (UniProt assession P48039): CHEMBL15060, agomelatine (CHEMBL10878), and iodomelatonin (CHEMBL289233). Most of the identified attributed atoms are positioned in close contact with the protein, indicating that the explanation methods emphasized ligand atoms likely to interact directly with the receptor, as seen in Figure 4b.

**Figure 4.**
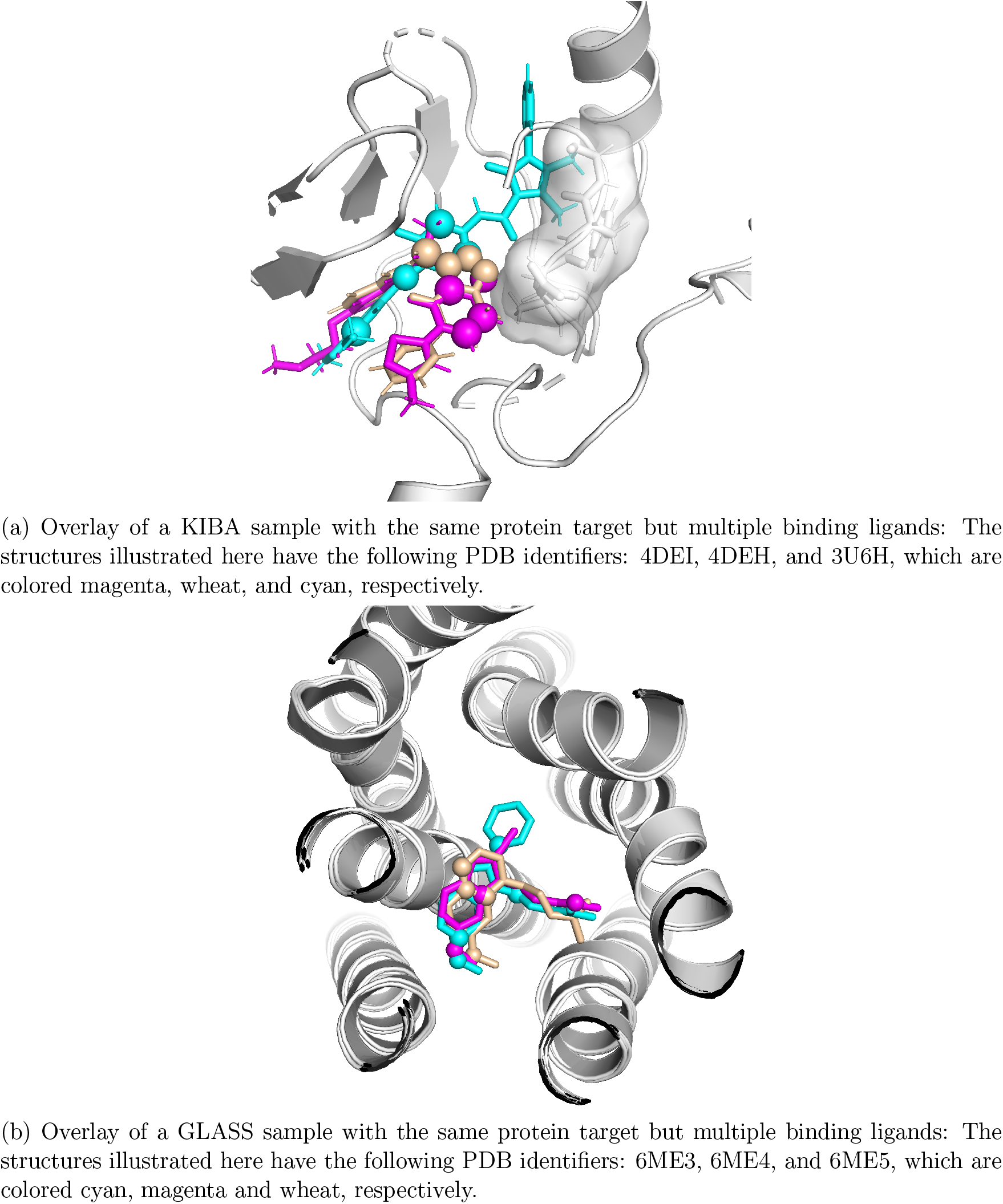
Example complexes from the KIBA and GLASS with multiple ligands. The three-dimensional structures were overlaid based on aligning the protein molecules. Attributed atoms are shown as balls in the stick representation of the ligands and colored in wheat, magenta, and cyan, respectively. The DFG regions in kinases in plot (a) are shown as the surface representation.

To further assess the spatial consistency of attributions for different ligands bound to the same protein, we compared the pairwise distances between the centers of mass (COM) of consensus-attributed atoms against the pairwise distances between the COM of all ligand atoms (Figure 5) from individual ligands. This analysis revealed that the attributed atoms from different ligands are not necessarily co-located in proximity to one another. Using the pairwise distance between the COM of all ligand atoms as a benchmark, the distances between the COMs of attributed atoms were consistently larger, indicating a lack of spatial convergence. We further analyzed the position of attributed atoms within individual ligands by measuring the distance from the COM of attributed atoms to the ligand’s total molecular COM, comparing it to the same measure for non-attributed atoms.6 The results indicate that consensus-attributed atoms are not centrally located but are instead enriched on the molecular periphery, consistent with their role at the protein-ligand interface. While the use of the COM has limitations—as it can be sensitive to the inclusion or exclusion of individual atoms in small sets—the overall trend is clear: attributed atoms across different ligands binding the same target are not necessarily co-located in a single, conserved spatial locus, but are consistently positioned at the ligand’s interaction surface.

**Figure 5.**
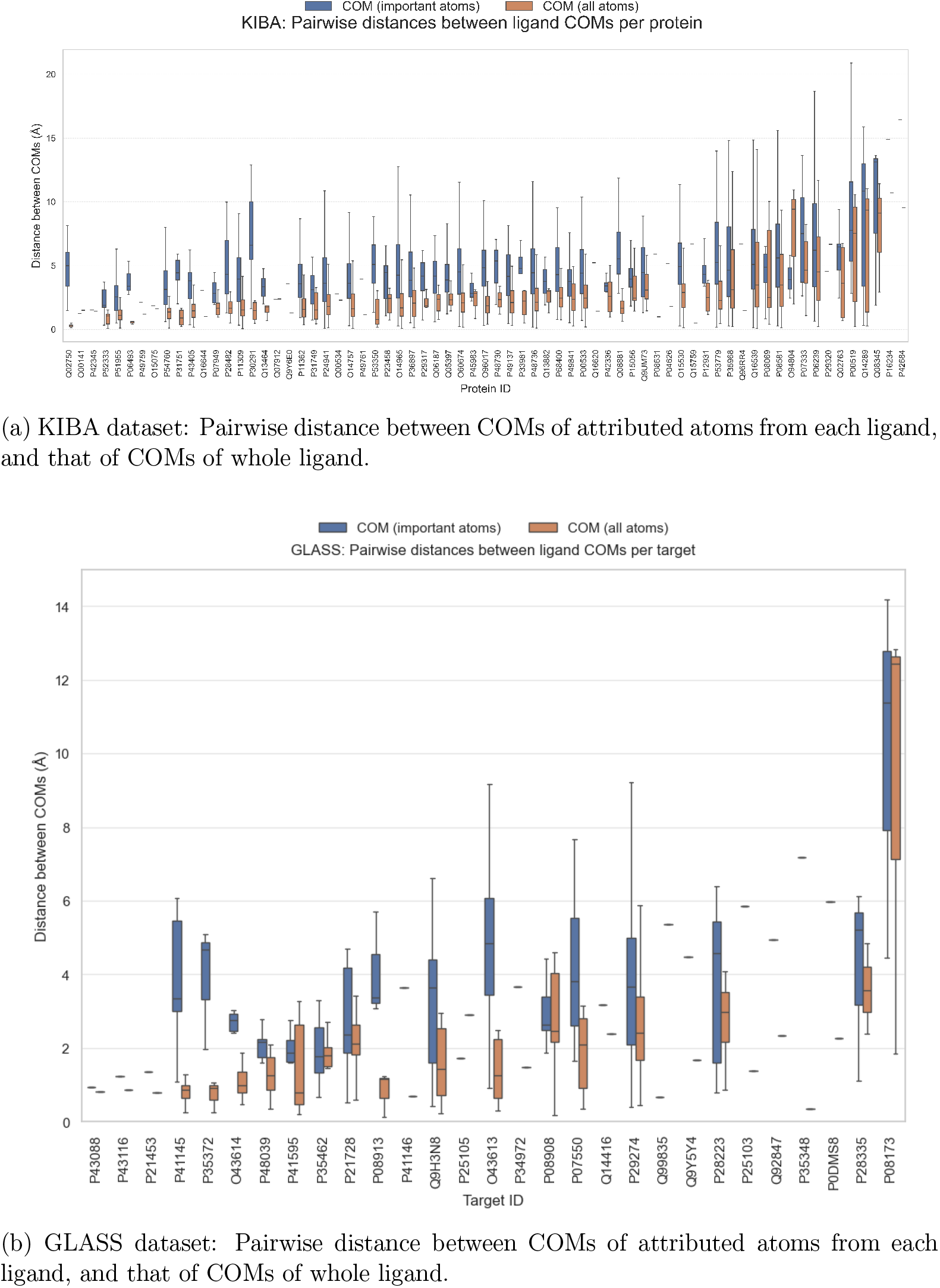
Comparison of pairwise center-of-mass (COM) distances for KIBA (a) and GLASS (b) datasets. The distances of attributed atoms are plotted in blue and the distances of all ligands’ atoms are colored in orange. Means and standard deviations for different ligands binding the same protein are shown.

**Figure 6.**
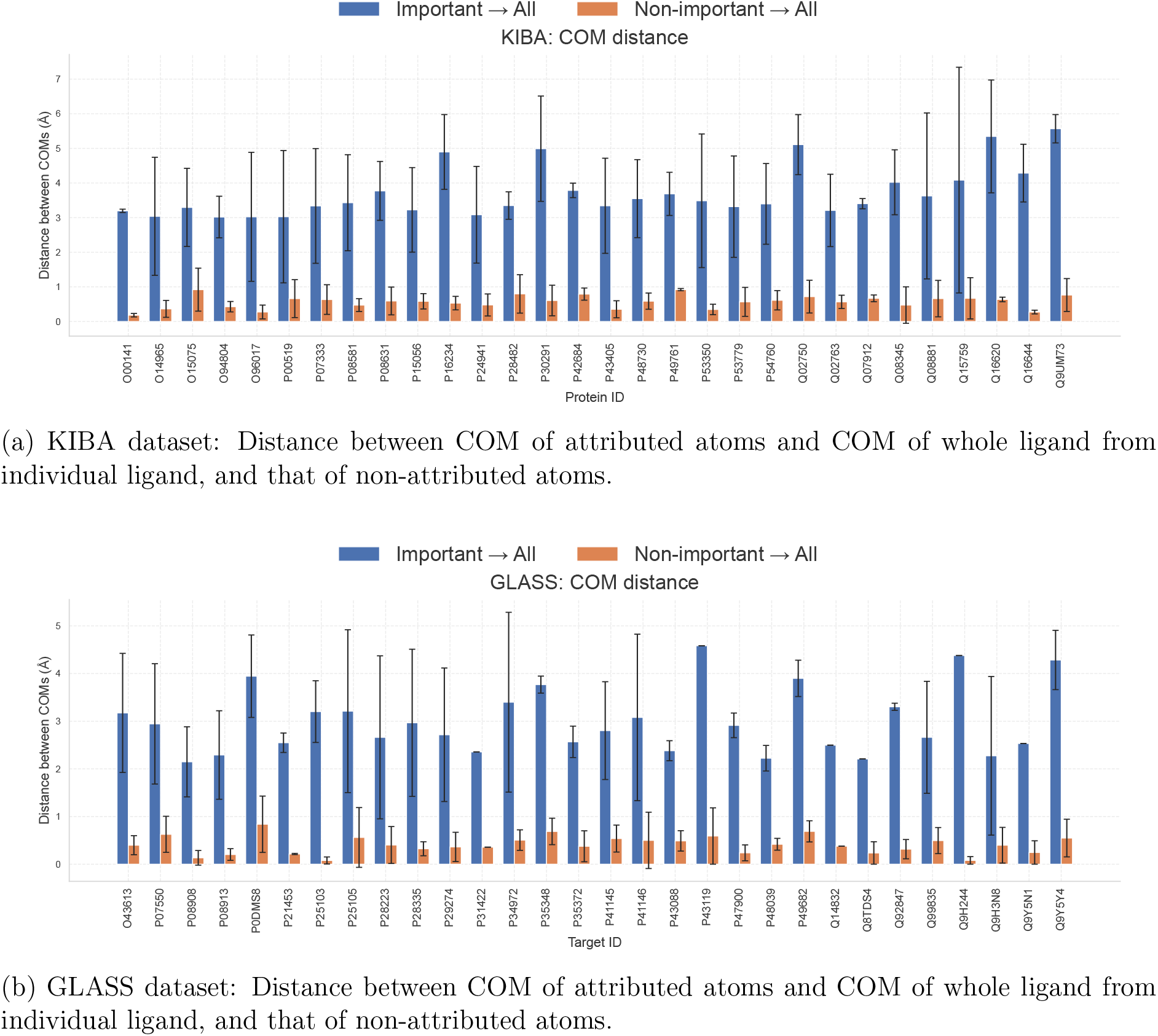
Comparison of center-of-mass (COM) distances between attributed atoms and whole ligands, vs. non-attributed atoms and whole ligands, for KIBA (a) and GLASS (b) datasets. The distances of attributed atoms are plotted in blue and non-attributed atoms are plotted in orange. Means and standard deviations for different ligands binding the same protein are shown.

## Conclusion

In this study, we investigated the explainability of graph neural network-based drug-target interaction models by applying four attribution techniques: Input x Gradient, Guided Back-propagation, Integrated Gradients and Gradient SHAP on two benchmark datasets, protein kinases and their inhibitors (KIBA) and G-protein coupled receptors and their ligands (GLASS). Our analysis revealed that the different methods often converged on overlapping subsets of ligand atoms, suggesting a degree of robustness across explanation techniques. These atoms also tend to directly contact the corresponding proteins, suggesting that they indeed play an important role in protein-ligand interactions. Moreover, when ligands binding to the same protein were overlaid, the attributed atoms, although located in the protein-contacting parts of the ligands, often contact different sites in the binding pocket, indicating the importance of different pharmacophores.

Overall, our results suggest that *post hoc* attribution methods can provide chemically meaningful insights into model predictions, offering a step towards interpretable and reliable applications of deep learning in drug discovery. Future work could expand on these findings by exploring additional explanation families and extending the analysis to larger and more diverse datasets.

## Acknowledgement

Q.L. would like to thank the Neuro-explicit Drug Discovery (NEDD) project of the Saarland University for financial support. O.V.K. acknowledges financial support from the Klaus Faber Foundation. We are grateful to Roman Joeres and Dr. Christiane Ehrt for a critical reading and discussion of the manuscript.

## Supporting Information Available

### Supplementary File S1 — KIBA structural dataset

File:`final_kiba.csv`

This file lists all protein–ligand complexes from the KIBA dataset that were used for the proximity analysis. Each entry includes the protein target identifier, drug (ligand) identifier, and the corresponding structure ID of the co-crystallised complex. These data define the subset of experimentally resolved structures used to evaluate the spatial correspondence between model-attributed atoms and binding pocket residues.

### Supplementary File S2 — GLASS structural dataset

File:`final_glass.csv`

This file provides analogous information for the GLASS dataset, including protein target identifiers, ligand identifiers, and complex structure IDs used for the proximity analysis. Together, these files document the structural basis of the datasets employed in the biological relevance evaluation.

### KIBA Score

The KIBA score integrates heterogeneous kinase inhibitor bioactivity measurements (IC_50_, *K*_*i*_, and *K*_*d*_) into a single unified metric. To harmonise the different assay formats, a model-based adjustment was introduced to rescale *K*_*i*_ and *K*_*d*_ using IC_50_, thereby increasing the consistency between bioactivity types.^4^

The adjustment formulas are:

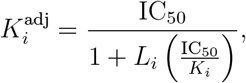

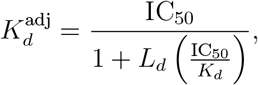

where *L*_*i*_ and *L*_*d*_ are the parameters that determine the weights of IC_50_ in the model-based adjustments for *K*_*i*_ and *K*_*d*_. The final KIBA score for each drug–target pair is defined as:

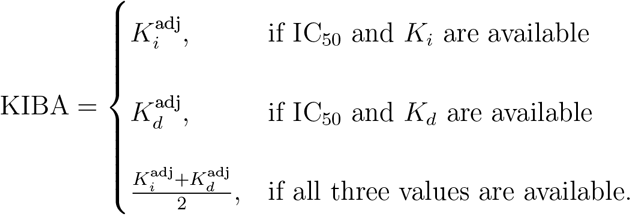

This integrated score provides a harmonised quantitative representation of the inhibitor bioactivity and improves the consistency between different assay types.

